# An Open-Source End-to-End Pipeline for Large-Scale EEG-Based Brain Age Modelling

**DOI:** 10.64898/2026.08.04.742735

**Authors:** Siddharth Rajesh, Deepshik Sharma, Rahul Venugopal, Arun Sasidharan, Saketh Malipeddi, Praerna Chowdhury, P. N. Ravindra

## Abstract

Aging affects individuals at varying biological rates, prompting the development of the Brain Age Index (BAI) to quantify neurobiological health relative to chronological age and disease risk. While structural MRI has dominated brain age prediction, its high cost, immobility, and low temporal resolution restrict its clinical scalability and responsiveness to transient neurophysiological changes. Electroencephalography (EEG) offers a highly scalable, portable, and temporally precise alternative capable of capturing dynamic brain states. However, the transition of EEG-based models to clinical biomarkers is impeded by methodological limitations, including small or biased datasets, inconsistent preprocessing pipelines, and a distinct lack of interpretable machine learning approaches. To address these persistent challenges, this paper presents a comprehensive, open-source, end-to-end pipeline for large-scale EEG-based brain age modeling. Developed using the Temple University Hospital EEG Corpus (TUEG) the largest publicly available resting-state EEG dataset. The pipeline encompasses rigorous data engineering, reproducible preprocessing, and robust feature extraction. Following quality control and subject-level dataset partitioning to definitively prevent data leakage, exactly 41,181 recordings were successfully retained. Two independent feature sets were extracted: the Catch22 time-series characteristics and a comprehensive set of spectral, aperiodic, and non-linear dynamics from the CCS toolbox. The methodology evaluates seven regression models, optimized via Optuna for hyperparameter tuning, and integrates SHAP (SHapley Additive exPlanations) for transparent feature importance analysis. By making this infrastructure publicly available, this work lowers the barrier to entry for large-cohort studies, fostering reproducible development and clinical validation of dynamic brain age biomarkers.

## Introduction

Aging is a complex biological process that is still not fully understood. What is clear, however, is that as we age, there is an increased risk of illness and mortality (Niccoli & Partridge, 2012). It is equally clear that humans do not age at the same biological rate, and that the onset and progression of age-related disease vary substantially across individuals (Cole et al., 2019). For this reason, considerable effort has gone into defining biological age as distinct from chronological age. Biomarkers such as telomere length, epigenetic clocks, and grip strength have been developed to capture the physiological burden of aging more accurately than chronological age alone, with the overarching goal of better predicting disease risk and residual lifespan (Cole et al., 2019).

Aging also affects the brain, and this motivated the development of biomarkers that quantify brain aging specifically. Early work hypothesised that neurodegenerative diseases reflect a form of accelerated brain atrophy, and that such pathological change might be detectable before clinical symptoms emerge (Franke et al., 2010). This led to the idea that an individual’s age could be estimated directly from brain anatomy, and more broadly, that machine learning models trained on neuroimaging data from healthy individuals across a wide age range could learn the normative trajectory of brain aging. Once trained, these models can be applied to clinical cohorts: if a participant’s brain shows greater structural decline than expected for their age, the model predicts an age higher than their true chronological age.

Since then, a substantial literature has developed around predicting age from neuroimaging data (Cole et al., 2017; Dosenbach et al., 2010; Franke et al., 2010, 2012; Jonsson et al., 2019). In the standard pipeline, high-dimensional MRI data, typically T1-weighted scans from healthy participants across a wide age range, are used to learn the normative aging trajectory. For instance, a relevance vector machine model estimated the brain age of Alzheimer’s disease patients to be, on average, 10 years above their chronological age (Franke et al., 2010). Critically, this overprediction is not a modelling artefact. It has been replicated consistently across independent studies and follows a graded pattern aligned with disease severity.

In this literature, the model-predicted age is termed Brain Age (BA), while the true age is Chronological Age (CA). The difference between the two is referred to here as the Brain Age Index (BAI) (BAI = BA – CA), a quantity that also appears under several related names including Brain Age Gap (BAG) (Zhang et al., 2025), Brain Age Delta (Smith et al., 2019), Brain Age Gap Estimation (BrainAGE) (Franke et al., 2010), and brain-predicted age difference (Boyle et al., 2020). Throughout this paper, we use the Brain Age Index (BAI) consistently. A positive BAI, where the brain appears older than its chronological age, is interpreted as a marker of accelerated neurobiological aging and has been associated with elevated risk for neurodegenerative disease, cognitive decline, and mortality. Conversely, a negative BAI, where the brain appears younger than its chronological age, is interpreted as a marker of decelerated neurobiological aging and has been associated with better cognitive outcomes, and resilience against age-related decline (Luders et al., 2016). This is particularly relevant for identifying activities with neuroprotective value like education and music that may delay the onset of aging-related disease (Rogenmoser et al., 2018; Steffener et al., 2016). Across the literature, model performance is standardly evaluated using Mean Absolute Error (MAE) and R² (de et al., 2022).

Beyond reflecting current disease state, BAI demonstrates genuine prognostic potential by predicting future neurological decline before the onset of overt clinical impairment. Evidence across the Alzheimer’s disease continuum illustrates this pattern: among individuals with mild cognitive impairment (MCI), patients who rapidly converted to Alzheimer’s disease exhibited a BAI of +8.73 years, compared to +5.62 years in late converters and only +0.75 years in stable MCI patients, suggesting that elevated BAI tracks not only disease presence but also progression rate (Gaser et al., 2013). Similar findings have been reported in psychosis research, where individuals classified as At Risk Mental State (ARMS) showed a BAI of +1.7 years prior to formal diagnosis, suggesting that accelerated brain aging may already be detectable during pre-clinical stages (Koutsouleris et al., 2014). The predictive value of BAI also extends to mortality outcomes. In a sleep EEG study involving 4,877 community-dwelling adults aged 40 years and older, higher BAI was associated with reduced life expectancy, with each standard deviation increase corresponding to a 12% greater hazard of all-cause mortality and a reduction of 0.81 years in expected lifespan (Paixao et al., 2020). Likewise, a longitudinal study of 2,609 older men without baseline dementia reported that individuals in the highest BAI quintile were 1.66 times more likely to develop cognitive impairment over an 11-year follow-up period than those in the lowest quintile, with brain age outperforming chronological age as a predictor of future cognitive decline (J. Sun et al., 2024). Collectively, these findings suggest that BAI functions not only as a cross-sectional marker of existing pathology, but also as a prospective biomarker capable of identifying elevated neurological risk before symptoms become clinically apparent.

Structural MRI has dominated brain age research and generally achieves strong predictive accuracy, with MAEs of 1.0 to 1.9 years in childhood and adolescence (ages 3–20) and 4.3 to 13.5 years across early to late adulthood (ages 19–86) (Franke & Gaser, 2019). However, it carries significant practical limitations. MRI scanners are expensive, immobile, and computationally demanding, restricting access largely to well-resourced institutions and introducing potential selection bias (Engemann et al., 2022). MRI also requires participants to remain still in a noisy, enclosed environment for extended periods, which is often infeasible for infants, older adults, and individuals with severe cognitive or psychiatric impairment (Bouma, 2024). Beyond these logistical constraints, MRI’s poor temporal resolution presents a more fundamental problem. Evidence has shown that brain age is not a static property: it is sensitive to transient experiential states, shifting measurably in response to creative activities. A study using M/EEG functional connectivity with machine learning across 1,240 participants found that expert practitioners in dance, music, visual arts, and strategy video games showed systematically delayed brain age compared to non-experts, with effects scaling with level of expertise (Coronel-Oliveros et al., 2025). Crucially, even short-term creative learning produced measurable reductions in BAI within the same individual. For instance, participants who engaged in 30 hours of video game training spread over 3 to 4 weeks showed a BAI of −3.1 years compared to pre-training baseline (Coronel-Oliveros et al., 2025).The sensitivity of brain age to transient biological states extends beyond behaviour: a study tracking BAI fluctuations across the menstrual cycle found that BAI reached −1.3 years during ovulation and correlated negatively with estradiol levels, suggesting that hormonal variation can also cause shift in brain age estimates meaningfully (Franke et al., 2015). These effects were mechanistically linked to plasticity-driven increases in brain network efficiency and biophysical coupling, particularly in frontoparietal regions known to be vulnerable to aging (Moguilner et al., 2024). Brain age is therefore a dynamic index that responds to experience on timescales far shorter than structural atrophy, timescales that MRI is ill-equipped to resolve. A modality with sufficient temporal sensitivity is needed if brain age is to function as a robust and responsive biomarker of brain health.

Electroencephalography (EEG) provides such an alternative. It is highly scalable, relatively inexpensive, and portable, and can be deployed across diverse settings including clinics, ambulatory environments, and at-home assessments. Unlike MRI, which captures slow macroscopic changes such as gray matter atrophy and ventricular enlargement (Cole et al., 2019; Hua et al., 2024), EEG operates at millisecond-level temporal resolution and can directly measure the rapid dynamics of oscillatory rhythms, and functional network connectivity (Engemann et al., 2022). Resting-state EEG is particularly attractive for brain age estimation because it reflects intrinsic baseline brain activity rather than task-driven responses, making it a strong candidate for deriving generalised markers of functional maturity and degeneration (Zhulduzbayev et al., 2026). Importantly, electrophysiological changes in connectivity and firing patterns may emerge earlier than overt structural atrophy, suggesting EEG could detect neurodegenerative risk at a stage when intervention remains possible (Al-Ezzi et al., 2024). For these reasons, EEG represents the most promising modality for building a dynamic, accessible, and temporally sensitive brain age biomarker.

Sleep EEG is the most commonly used data in Brain Age Index research for good reasons: sleep EEG captures a full repertoire of distinct neural states (Wake, N1, N2, N3, and REM), reduces the influence of external confounds such as attention lapses and task non-compliance, and is less susceptible to the transient cognitive and emotional fluctuations that can introduce noise (Banks et al., 2025; H. Sun et al., 2019; J. Sun et al., 2024; Ye et al., 2020). Sleep-based models perform best, with MAEs as low as 4.19 years, reflecting the richness of sleep architecture as a neurobiological marker (Zhulduzbayev et al., 2026). However, resting-state EEG has been widely used to estimate brain age precisely because of its practical accessibility, no specialised sleep laboratory, no overnight recording, and no staging expertise, making it far more scalable for large-cohort research. A proof-of-concept study using a stacking ensemble on 468 participants established the feasibility of this approach, achieving an R² of 0.37 and an MAE of 6.87 years, modest performance that nonetheless confirmed resting-state EEG carries meaningful age-related information, while identifying reliable feature extraction as the central methodological challenge (Al Zoubi et al., 2018).

In classical machine learning pipelines for resting-state EEG, researchers typically extract spectral power features, entropy-based descriptors, and statistical moments such as skewness and kurtosis. While these features are selected based on prior neurophysiological knowledge, a model that predicts age accurately does not necessarily reveal which features are doing the predictive work nor whether those features reflect genuine biological aging or dataset-specific noise. This distinction matters enormously in brain age research, where the ultimate goal is not prediction alone, rather, identifying the EEG signatures that track neurobiological aging could illuminate the processes driving cognitive decline and provide the biological grounding necessary for clinical adoption. It is here that explainable AI (XAI) becomes indispensable, providing tools to probe trained models, attribute predictive weight to specific features, and determine whether a model’s decisions align with known neuroscience rather than incidental statistical regularities (Lyberatos et al., 2026). For a more detailed overview of the feature sets used in the EEG based Brain Age literature, refer to (Zhulduzbayev et al., 2026). An extension of this approach, the Chronnectomic Brain Age Index (CBAI), uses dynamic functional connectivity and phase synchronisation to capture temporal evolution in brain network organisation as a maturation index (Dimitriadis & Salis, 2017). A recent benchmarking study across four international M/EEG datasets with over 2,500 participants found that spatially aware representations achieved the strongest performance, while handcrafted features combined with random forest regression remained a robust baseline across varied conditions (Engemann et al., 2022; Iyer et al., 2023). In terms of overall performance, current models achieve R² values between 0.60 and 0.74 in adult populations, with MAEs in the range of 4 to 8 years. At the other end of the lifespan, functional brain age models in neonates are considerably more precise, with MAEs of approximately 1 to 1.3 months (An et al., 2025; Davoudi et al., 2025; Zhulduzbayev et al., 2026). This marked improvement in predictive accuracy reflects the non-linearity of brain aging: during early development (0-5 years), the brain undergoes rapid and pronounced structural change, providing machine learning models with strong, easily learnable signals. As the rate of change decelerates with advancing age, the features distinguishing one year from the next become increasingly subtle, demanding far greater data to learn reliably. The high accuracy in neonatal models is therefore less a reflection of superior methodology and more a consequence of the biology itself (Bethlehem et al., 2022).

Despite these promising results, the field faces persistent methodological limitations that collectively impede the transition of EEG-based brain age estimation from a research signal to a validated clinical biomarker. Most models are evaluated only through internal cross-validation on a single dataset, increasing the risk of overfitting and limiting generalisability(An et al., 2025; Davoudi et al., 2025; Dimitriadis & Salis, 2017). Many datasets are also demographically imbalanced, with heavy representation of young and middle-aged adults and poor coverage of children, older adults, and populations outside high-income countries (Engemann et al., 2022).

Fewer than a quarter of studies apply bias correction procedures such as linear residualisation, leaving indices like BAI susceptible to regression-to-the-mean effects in which younger subjects are systematically over-predicted and older subjects under-predicted, producing estimates that are statistically distorted and clinically difficult to interpret (Zhulduzbayev et al., 2026). At the methodological level, preprocessing pipelines vary widely in filter settings, artifact rejection strategies, epoch lengths, and electrode montages, making cross-study comparison extremely difficult (Engemann et al., 2022). Important biological and clinical confounders including medication use, sleep quality, sex, and recording conditions such as eyes-open versus eyes-closed are often inconsistently controlled or unreported (Zhulduzbayev et al., 2026). Finally, while deep learning is increasingly adopted, interpretability analyses remain uncommon, leaving the neurobiological basis of model predictions unclear. Given that most available datasets contain only hundreds to low thousands of participants, deep learning has also not demonstrated a consistent advantage over classical machine learning in this domain (Ettling et al., 2024).

Taken together, these limitations point to a clear need for a scalable, transparent, and reproducible infrastructure for large-scale EEG-based brain age research. To address this, we present an open-source, end-to-end pipeline developed on the Temple University Hospital EEG Corpus (TUEG), the largest publicly available resting-state EEG dataset, comprising 69,670 EDF files (Obeid & Picone, 2016). The pipeline covers the full analytical workflow, from large-scale data engineering and reproducible preprocessing through feature extraction and systematic feature importance analysis. By making this publicly available, we aim to lower the barrier to large-cohort EEG research and provide a reproducible foundation for developing, comparing, and validating future brain age models.

## Methodology

### Dataset

The Temple University Hospital EEG Corpus, specifically the TUEG dataset, version 2.0.1, was used in this study. The dataset comprises clinical EEG recordings collected from archival records at Temple University Hospital (TUH), Philadelphia, USA, spanning 14,987 patients and 69,670 EDF files. Considerable heterogeneity exists across recordings with respect to sampling rate (87% of files were sampled at 250 Hz, 8.3% of files sampled at 256 Hz, 3.8% of files sampled at 400 Hz, 1% of files sampled at 512 Hz), channel configuration (Unipolar montages with Average Reference and Linked Ears), and recording duration (min = 1s, max = 88425s). Access to the dataset was granted following approval by the TUH-EEG data governance team and was obtained via secure SSH transfer. The dataset download structure contains a headers text file, a readme, and the corresponding EDF recordings.

### Data Engineering

All data engineering operations were performed exclusively on the metadata contained within the headers text file, ensuring no EEG signal data were read, modified, or distorted at this stage. At each filtering step, a CSV file was generated retaining a filepath column as a constant identifier, which was later used to call the relevant EDF files during pre-processing and feature extraction.

The TUEG dataset includes recordings in two referencing configurations: average reference and linked-ears reference. Given that the average reference configuration afforded the greatest file coverage, only recordings in this configuration were retained. Within this subset, there was heterogeneity in the number of edf files that contained all the electrodes. Only 16 of the following channels: Fp1, Fp2, F3, F4, C3, C4, P3, P4, O1, O2, F7, F8, T3, T4, T5, and T6, were included in majority of the edf files and remained consistent throughout as identified using the script 00_get_summary_from_txt.py and filtered using 01_selecting_16chans.py. This yielded an initial working pool of 56,848 EDF files.

Files were subsequently screened for demographic completeness using 02_valid_age_gender_files.py. Recordings with missing age, missing gender, or an age value of 999 (a known sentinel value for missing data in the corpus) were excluded. Age distribution was plotted (see Figure 4). There were a total of 19819 males and 21362 females in the final dataset. Since, the number of people below the age of 18 and above the age of 90 were significantly low, we removed them to ensure adequate sample size for the ML models to learn.

**Figure 1:**
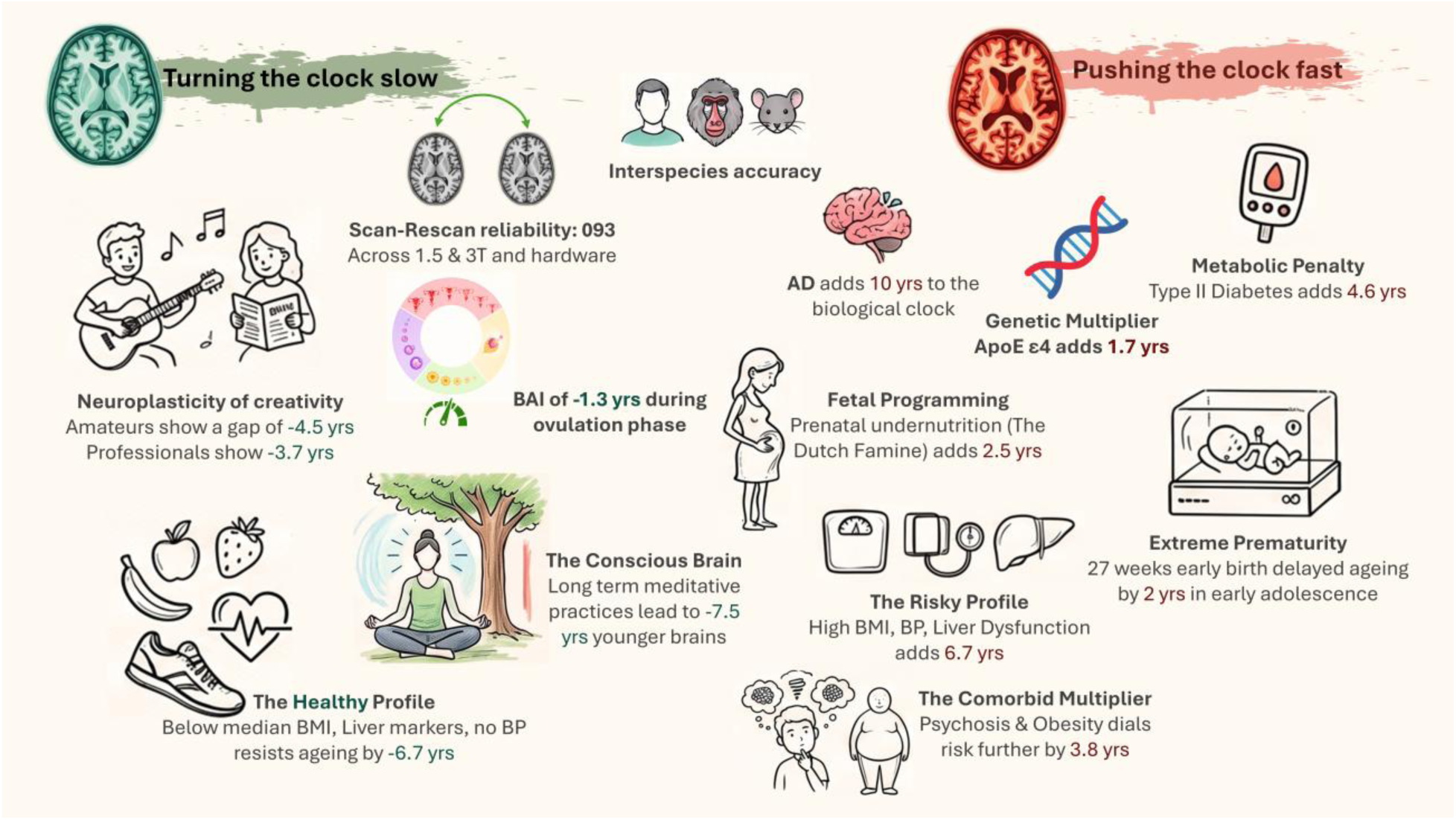
Model Accuracy and Versatility: The BrainAGE model demonstrates high accuracy across species (human adults and children, baboons, and rodents) and is highly stable scan-rescan reliability (ICC) of 0.93 across different hardware. Short-term Biological Sensitivity: The model can capture transient neurological fluctuations, such as a -1.3-year decrease in biological brain age during ovulation driven by higher estradiol levels. Predicting Alzheimer’s Disease (AD): BAI predicts the conversion from Mild Cognitive Impairment (MCI) to AD within 12 months with 81% accuracy, outperforming gold-standard biomarkers like hippocampus volume and cognitive scores. Neurodegenerative Acceleration: A baseline BAI above the median indicates a nearly 4x greater risk of AD conversion, and patients in the 4th quartile face a 4.7x greater risk. Every additional year in BAI compounds the risk of AD conversion by 10%. Disease Trajectories: AD fundamentally accelerates the biological clock; at diagnosis, patients exhibit a +10-year baseline gap, and their brains age 1.5 biological years for every chronological year. Genetic Multipliers: The APOE ε4 gene acts as a longitudinal accelerator, adding up to 1.7 years of brain aging per year in AD patients. Metabolic Health Toll: Systemic health directly dictates neuroanatomical aging. Type 2 Diabetes imposes an immediate +4.6-year penalty to brain age and continues to add +0.2 years for every subsequent chronological year. Compounding Health Risks: In cognitively healthy older men, metabolic factors like BMI, blood pressure, and liver function account for 39% of the variance in brain aging. This creates a nearly 15-year swing between "healthy" profiles (-8.0 years) and "risky" profiles (+6.7 years). Psychiatric Divergence: First-episode schizophrenia accelerates brain aging by +2.6 years. When compounded with obesity as a comorbid risk factor, this penalty jumps to +3.8 years. Bipolar disorder, however, shows no significant deviation from healthy aging. Fetal and Early Life Programming: Biological brain age is influenced decades before clinical symptoms appear. Men exposed to severe prenatal undernutrition (like the Dutch Famine) show a permanent +2.5-year brain aging penalty that persists into their late 60s. Extreme preterm birth (<27 weeks) causes delayed structural maturation, yielding a -2.0-year BAI in early adolescence. Protective Lifestyle Decelerators: Long-term meditation actively protects against structural atrophy, resulting in brains estimated to be 7.5 years younger at age 50. For every chronological year past age 50, a long-term meditator’s brain ages approximately 1 month and 22 days slower than a control subject. Neuroplasticity of Creativity: Amateur musicians show a -4.5-year brain age reduction, though professional musicians show a slightly smaller benefit (-3.7 years), suggesting that performance pressure and occupational stress may dampen the protective effects.

**Figure 2:**
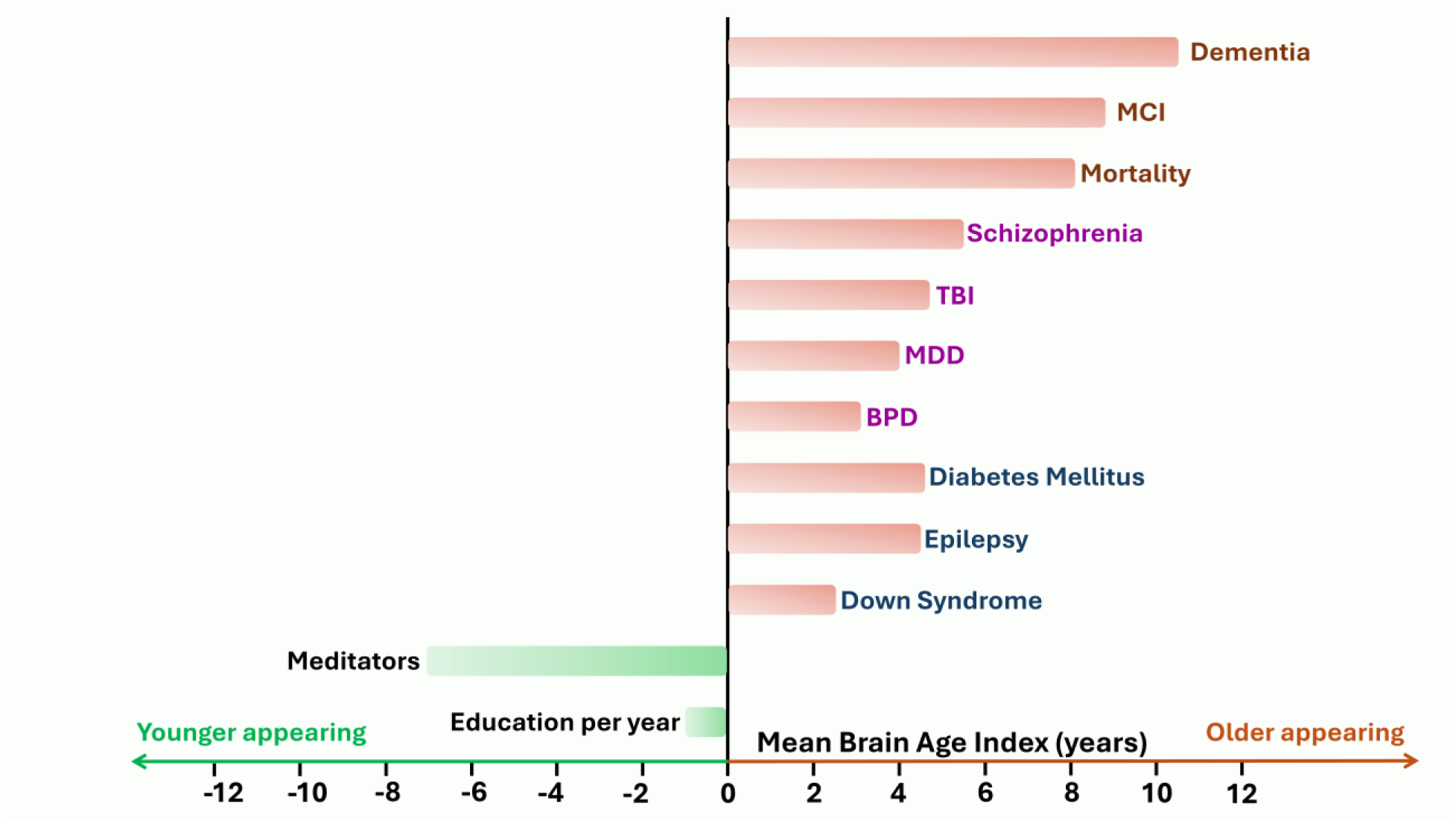
Brain Age Index (BAI) across clinical and protective conditions. Horizontal bars represent the range of BAI values reported across independent studies for each condition. BAI = 0 represents the chronological age. Positive BAI values indicate accelerated brain aging relative to chronological age, while negative BAI values reflect decelerated aging. MCI - Mild Cognitive Impairment, TBI - Traumatic Brain Injury, MDD - Major Depressive Disorder, BPD - Borderline Personality Disorder.

**Figure 3:**
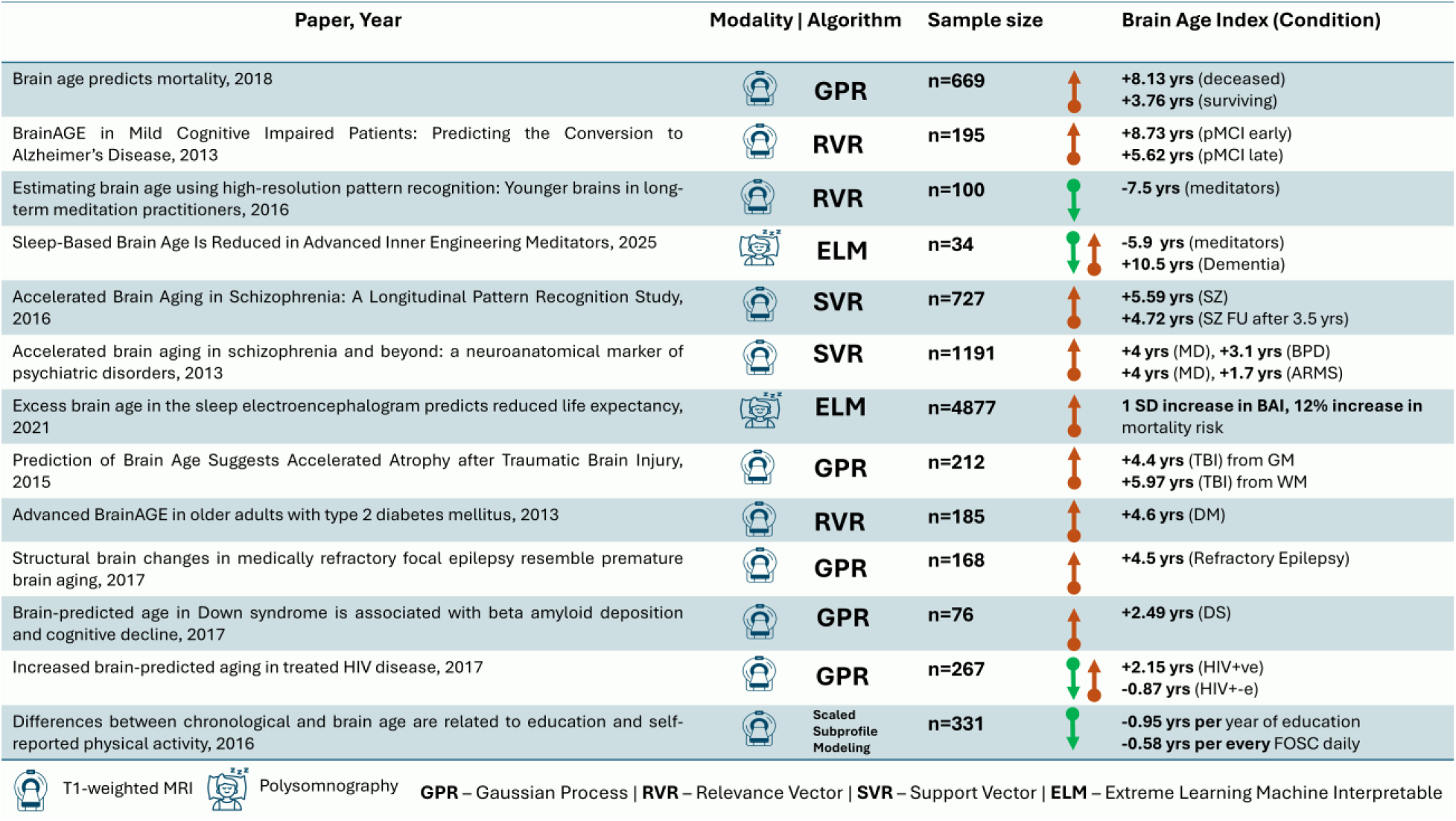
Overview of Brain Age Index (BAI) scores across pathological and neuroprotective conditions reported in representative studies. The table highlights the imaging modality, machine learning model, cohort size, and reported brain age differences relative to healthy controls. Positive values indicate accelerated brain ageing, whereas negative values indicate younger brain age. pMCI, progressive mild cognitive impairment; SZ, schizophrenia; MD, major depression; BPD, borderline personality disorder; ARMS, at-risk mental state; TBI, traumatic brain injury; DM, diabetes mellitus; DS, Down syndrome; HIV, human immunodeficiency virus; FOSC, flights of stairs climbed.

**Figure 4:**
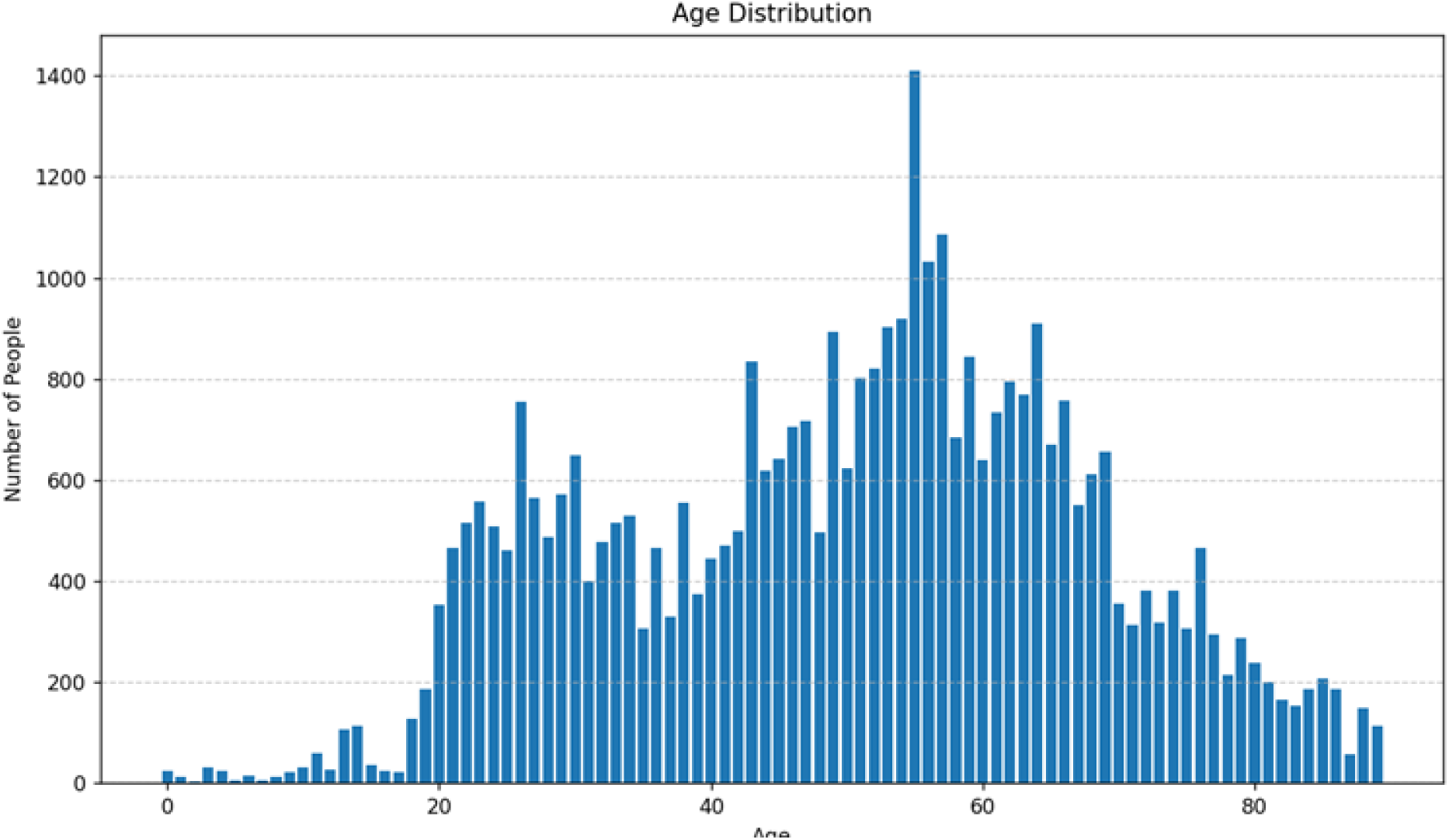
Age distribution of all the valid files in TUEG dataset.

Because TUEG contains multiple recording sessions per subject, including a small number of longitudinal recordings in which the same subject appears at different ages, dataset partitioning was performed at the subject level to prevent data leakage across splits (Narayanan & Kapoor, 2024). Unique subjects were identified using 03_unique_subjects.py, and a stratified split of 70% training, 15% validation, and 15% test was manually done at the subject level. The edf files for the subjects were repopulated using 04_generating_splits.py. A dedicated sanity check script (sanity_check_for_data_split.py) was run to verify that no subject appeared in more than one split. After all filtering and partitioning steps, the final dataset comprised 41,181 EDF files: 28,827 in the training set, 6,177 in the validation set, and 6,177 in the test set. Each split was stored as a CSV with columns for filepath, age, and gender.

### Pre-processing

All pre-processing was implemented in Python using MNE-Python (Gramfort et al., 2014). For each EDF file, a high-pass filter at 0.5 Hz was applied first, followed by a low-pass filter at 45 Hz, each using MNE’s default filter settings. A notch filter at 60 Hz was then applied to suppress power-line interference. Signals were subsequently resampled to 200 Hz to standardise sampling rates across the dataset. The continuous filtered and resampled signal was then segmented into non-overlapping 30-second epochs.

Artefact rejection was applied in two sequential levels. In the first level, a global amplitude threshold was applied: any epoch in which the signal amplitude exceeded ±500 µV in any channel was rejected outright. In the second level, more refined artefact detection was applied, with the strategy adapted according to the number of surviving epochs per file. For files with fewer than four epochs, they were rejected due to the low number of epochs and were excluded from further analysis. For files with between four and ten epochs, MNE’s get_rejection_threshold function was used to estimate a data-driven global rejection threshold, and epochs exceeding this threshold were discarded. For files with more than ten epochs, the Autoreject library was used, which estimates channel-wise peak-to-peak rejection thresholds and interpolation parameters in a cross-validated manner. Autoreject was run with n_interpolate = [1, 2, 3, 4] and consensus = [0.25]. Files for which all epochs were rejected following both levels of artefact rejection were excluded from further analysis.

For each retained file, features were subsequently computed across all clean epochs, with per-epoch values aggregated by computing the mean and standard deviation, yielding two summary statistics per feature per channel.

### Feature Extraction

Feature extraction was performed separately for the training, validation, and test sets to prevent any information leakage during model development. Two independent feature sets were derived from the pre-processed EEG data.

**Catch22 features *-*** Time-series features were extracted using the pycatch22 library, which implements the 22 canonical features of the CAnonical Time-series CHaracteristics (Catch22) set (Lubba et al., 2019).

For each file, features were extracted per channel, yielding 16 rows per file (one per channel). The mean and standard deviation of each feature across clean epochs were computed, resulting in 44 feature columns (22 features × 2 statistics). Additional columns included the filepath, age, one-hot encoded gender (2 columns), channel label, and the three-dimensional Cartesian electrode coordinates (x, y, z), for a total of 51 columns per row.

A second feature set was derived using a Python toolbox for EEG analysis developed by the Centre of Consciousness Studies, NIMHANS. This feature set comprises spectral, aperiodic, and non-linear EEG features (Sasidharan, 2026).

Power spectral density (PSD) was estimated using Welch’s method, and band-limited power was computed across canonical frequency bands (1–40 Hz). Aperiodic and oscillatory components were separated using FOOOF-like and IRASA-based approaches, from which band-wise oscillatory power and summary aperiodic parameters (e.g., slope and intercept) were extracted.

Non-linear temporal dynamics were quantified using measures of entropy (permutation entropy, sample entropy), fractal complexity (Petrosian, Katz, and Higuchi fractal dimensions), Lempel–Ziv complexity, detrended fluctuation analysis (DFA), and singular value decomposition (SVD) entropy. In addition, the autocorrelation window (ACW) was computed as the lag at which the autocorrelation function decayed below 0.5. Multifractal properties were assessed using MF-DFA applied to band-limited amplitude envelopes, yielding the generalized Hurst exponent, spectrum width, and peak location.

As with the Catch22 set, the mean and standard deviation across clean epochs were computed per feature, yielding 138 feature columns (69 × 2). The full CCS feature matrix thus contained 145 columns per row, including filepath, age, one-hot encoded gender, channel label, and electrode coordinates.

### Model Training and Evaluation

Seven regression models were trained and evaluated for the task of chronological age prediction from EEG features: Linear Regression (LR), Support Vector Regression (SVR), Relevance Vector Regression (RVR), Decision Tree (DT), Random Forest (RF), eXtreme Gradient Boosting (XGBoost), and Gaussian Process Regression (GPR). All models were implemented in Python; scikit-learn was used for LR, SVR, RVR, DT, and RF; XGBoost was implemented using the XGBoost library; and GPyTorch was used for GPR.

### Hyperparameter Optimization with Optuna

Hyperparameter optimization was performed using Optuna (Akiba et al., 2019) for all models where manual external tuning was required. For each trial, Optuna sampled a candidate configuration from a predefined search space, trained the model using only the training data, and evaluated the candidate using cross-validated mean absolute error (MAE). The best configuration was selected as the one that minimized the objective value. For tree-based and kernel-based models, the objective was estimated with 5-fold cross-validation on the training set; the held-out validation set was reserved for final reporting. After tuning, each model was retrained using the best hyperparameters and then evaluated on the validation and test sets.

A standard linear regression model from scikit-learn was used as a baseline. This model does not expose meaningful hyperparameters for optimization, and therefore no Optuna-based tuning was applied. The model was trained directly on the training data, minimizing mean squared error (MSE).

For the decision tree regressor, Optuna searched over max_depth, min_samples_leaf, min_samples_split, and ccp_alpha. The search was limited to a discrete space of depth values from 4 to 32, leaf sizes from 1 to 100, split thresholds from 2 to 200, and ccp_alpha values sampled on a logarithmic scale from (1e-5 to 1e-1). The number of trials was set to 200, and the objective minimized 5-fold cross-validated MAE.

For the random forest regressor, Optuna tuned n_estimators, max_depth, min_samples_leaf, min_samples_split, and max_features. The search space covered 200 to 800 trees, depths from 6 to 20, leaf sizes from 5 to 100, split thresholds from 5 to 100, and max_features sampled continuously between 0.3 and 1.0. The search used 200 trials and minimized 5-fold cross-validated MAE.

For XGBoost, the search space included n_estimators, max_depth, learning_rate, subsample, colsample_bytree, gamma, reg_alpha, and reg_lambda. The number of boosting rounds ranged from 200 to 800, tree depth from 3 to 10, learning rate from (1e-3 to 1e-1) on a logarithmic scale, subsampling ratios from 0.6 to 1.0, column sampling ratios from 0.4 to 1.0, split regularization gamma from 0.0 to 5.0, L1 regularization reg_alpha from (1e-5) to 1.0 on a logarithmic scale, and L2 regularization reg_lambda from (1e-2) to 10.0 on a logarithmic scale. This search was also run for 200 trials in the code version used here, with 5-fold cross-validated MAE as the objective.

For the support vector regressor, the kernel was fixed to RBF and only the most influential parameters were optimized: C, epsilon, and gamma. The search space used a logarithmic range for C from (1e-1) to (1e3), a uniform range for epsilon from 0.01 to 3.0, and a logarithmic range for gamma from (1e-5) to (1e-1). The SVR pipeline included standardization as part of the model so that scaling was performed within each cross-validation fold. The objective minimized 5-fold cross-validated MAE over 200 trials.

For relevance vector regression, the kernel was fixed to RBF and the search was restricted to gamma only, using a logarithmic range from (1e-5) to (1). This kept the optimization focused and avoided unnecessary search over solver controls that primarily affect numerical behavior rather than predictive performance. The objective again minimized 5-fold cross-validated MAE, with 200 trials.

Gaussian process regression with GPyTorch was not wrapped in Optuna, because the GP hyperparameters were already learned directly by maximizing the marginal log likelihood during training. In that case, the external optimization problem would be redundant. The model was trained with Adam on the negative marginal log likelihood, while the validation set was used only for monitoring.

Across all experiments, the final model for each method was retrained using the best hyperparameter configuration selected by Optuna, and predictions were subsequently evaluated on the validation and test sets using MAE, RMSE, and (R^2^).

### Feature Importance

To interpret model predictions, SHAP (SHapley Additive exPlanations) values (S. Lundberg & Lee, 2017) were computed on the held-out test set for all seven models. Model-appropriate SHAP estimators were selected to ensure both computational efficiency and methodological rigour. For linear regression, LinearSHAP was applied, which provides exact Shapley values by exploiting the closed-form structure of the linear model. For the tree-based models: Decision Tree, Random Forest, and XGBoost, TreeSHAP (S. M. Lundberg et al., 2020) was used, which computes exact Shapley values in polynomial time by traversing the decision tree structure directly. For the kernel-based and probabilistic models like Support Vector Regression, Relevance Vector Regression, and Gaussian Process Regression, KernelSHAP (S. M. Lundberg & Lee, 2017) was employed, a model-agnostic estimator that approximates Shapley values by sampling feature coalitions around a background reference distribution. As KernelSHAP is computationally expensive on high-dimensional datasets, the training background was compressed to 100 representative points using k-means summarisation prior to explanation, a standard practice for scaling KernelSHAP to datasets of this size. In all cases, SHAP values were computed using the training set as the background reference to establish the baseline expected prediction E[f(x)], while attributions were evaluated on the test set to reflect model behaviour on unseen data.

### Software and Reproducibility

The full analysis pipeline was implemented in Python. Key libraries used included MNE-Python (pre-processing) (Gramfort et al., 2014), Autoreject (artefact rejection) (Jas et al., 2017), pycatch22 (Catch22 features) (Lubba et al., 2019), scikit-learn (regression models) (Pedregosa et al., 2011), XGBoost (Chen & Guestrin, 2016), GPyTorch (Gardner et al., 2018), Optuna (hyperparameter optimisation) (Akiba et al., 2019), SHAP (feature importance) (S. M. Lundberg & Lee, 2017), and joblib (parallelisation). The code used in this study is openly available on GitHub at https://github.com/sidcodes37/Brain_Aging_CCS, and the library versions and key software dependencies required for reproducibility are documented in the repository.

## Notes

https://github.com/sidcodes37/Brain_Aging_CCS

### Competing Interest Statement

The authors have declared no competing interest.

