## Supplementary table for "An Open-Source End-to-End Pipeline for Large-Scale EEG-Based Brain Age Modelling"

### Supplementary Material

Age and Gender distribution is plotted across the three datasets below (see Figure 4 & Figure 5).

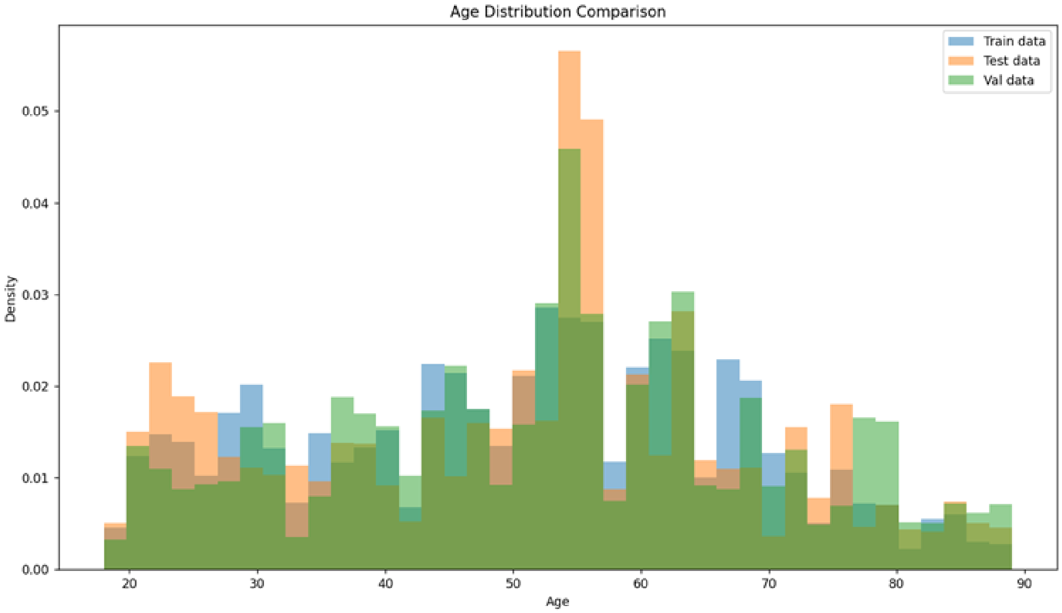


**Figure 5:** Age Distribution compared across train, test and val sets.

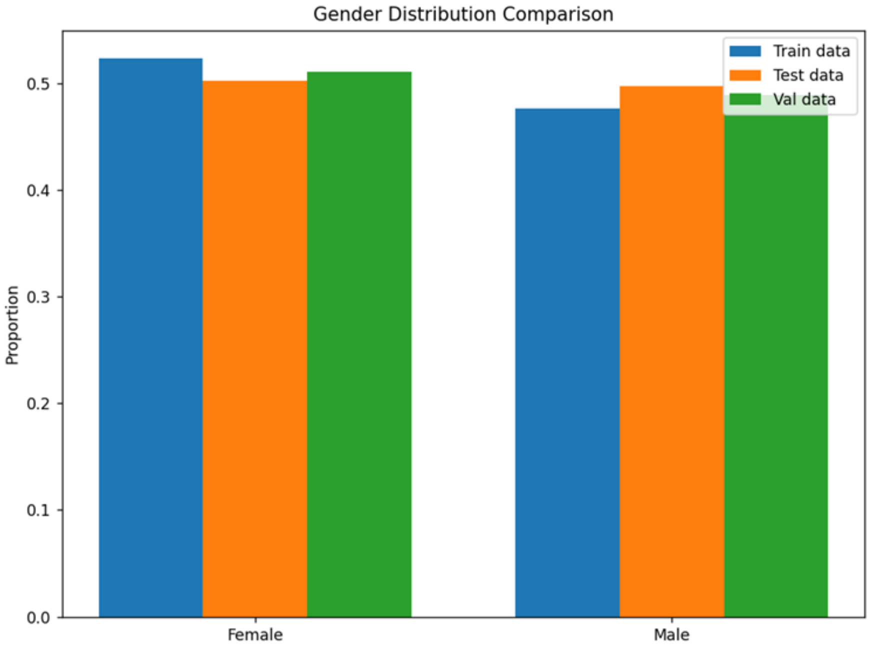


**Figure 6:** Gender Distribution compared across train, test and val sets.

*Predictive Value of BAI*

| Paper | Year | Modality | Test Dataset | ML Model | BAI |
|---|---|---|---|---|---|
| Brain age predicts mortality | 2018 | T1-weighted MRI | N = 669 Mean age = 72.67 years | GPR | BAI(deceased) = +8.13 years BAI(surviving) = +3.76 years |
| BrainAGE in Mild Cognitive Impaired Patients: Predicting the Conversion to Alzheimer's Disease | 2013 | T1-weighted MRI | N=195 with MCI at baseline | RVR | BAI(pMCI_early) = +8.73 years BAI(pMCI_late) = +5.62 years BAI(sMCI) = +0.75 years |
| Estimating brain age using high-resolution pattern recognition: Younger brains in long-term meditation practitioners | 2016 | T1-weighted MRI | N = 50 (long-term meditators) N = 50 (control) Mean age = 51.4 years | RVR | BAI(meditators) = -7.5 years |
| Sleep-Based Brain Age Is Reduced in Advanced Inner Engineering Meditators | 2025 | Sleep-based EEG | N = 34 Mean age = 38 yrs 36% female | Interpretable ML model (Adapted from Sun et al., 2019) | BAI (meditators) = -5.9 years BAI(MCI) = +8.8 years BAI(Dementia) = +10.5 years |
| Accelerated Brain Aging in Schizophrenia: A Longitudinal Pattern Recognition Study | 2016 | T1-weighted MRI | N = 341 (SZ patients) N = 386 (healthy population | SVR | BAI(SZ) = +5.59 years BAI(SZ FU after 3.48 yrs) = +4.72 years |

| | | | | | |
|---|---|---|---|---|---|
| | | | Baseline age = 16-67 years<br>Follow-up (FU) = 1-13 years | | |
| Accelerated brain aging in schizophrenia and beyond: a neuroanatomical marker of psychiatric disorders | 2013 | T1-weighted MRI | N=141 Schizophrenia (SZ) patients<br>N = 104 Major depression (MD)<br>N=57 BPD<br>N=89 ARMS<br>N= 800 healthy population (for training)<br>18-65 years | SVR | BAI(SZ) = +5.5 years<br>BAI(MD) = +4 years<br>BAI (BPD) = +3.1 years<br>BAI(ARMS) = +1.7 years |
| Excess brain age in the sleep electroencephalogram predicts reduced life expectancy | 2021 | Sleep-based EEG | N = 4877 healthy population<br>Age = 40+ | Extreme learning machine (ELM) | For every 1 SD increase in BAI, 12% increase in mortality risk & 0.81-year decrease in LE |
| Prediction of Brain Age Suggests Accelerated Atrophy after Traumatic Brain Injury | 2015 | T1-weighted MRI | N = 99 (TBI patients)<br>Mean age of TBI = 37.98±12.43 years<br>N = 113 | GPR | BAI(TBI) = +4.66 years for GM<br>BAI(TBI) = +5.97 years for WM |

| | | | | | |
|---|---|---|---|---|---|
| | | | (Healthy population)<br>Mean age of Healthy population = 43.3±20.24 years | | |
| Advanced BrainAGE in older adults with type 2 diabetes mellitus | 2013 | T1-weighted MRI | N = 98 (Type 2 DM)<br>N = 87 (Healhy population)<br><br>Mean age = 64.6 | RVR | BAI(DM) = +4.6 years |
| Structural brain changes in medically refractory focal epilepsy resemble premature brain aging | 2017 | T1-weighted MRI | N = 94 (medically refractory focal epilepsy)<br>N = 74 (healthy controls)<br>Age = 12-60 years | GPR | BAI(medically refractory epilepsy) = +4.5 years |
| Brain-predicted age in Down syndrome is associated with beta amyloid deposition and cognitive decline | 2017 | T1-weighted MRI | N = 46 (DS patients)<br>Mean age of DS = 42.30 ± 8.73 years<br>N = 30 (Healthy | GPR | BAI(DS) = +2.49 years |

| | | | | | |
|---|---|---|---|---|---|
| | | | population) Mean age of control = 46.23 ± 9.75 years | | |
| Increased brain-predicted aging in treated HIV disease | 2017 | T1-weighted MRI | N = 162 (HIV-positive) Age of HIV patients = 45-82 years N = 105 (HIV-negative) | GPR | BAI(HIV-positive) = +2.15 years BAI(HIV-negative) = -0.87 years |
| Differences between chronological and brain age are related to education and self-reported physical activity | 2016 | T1-weighted MRI | N = 331 (healthy population) Age = 19-79 years | Scaled subprofile modeling | BAI(Education) = -0.95 years/ year of education BAI(FOSC) = -0.58 years per every additional FOSC daily |
